# Influence of metal cations on the viscoelastic properties of *Escherichia coli* biofilms

**DOI:** 10.1101/2022.09.29.510089

**Authors:** Adrien Sarlet, Valentin Ruffine, Kerstin G. Blank, Cécile M. Bidan

## Abstract

Biofilms frequently cause complications in various areas of human life, e.g. in medicine and in the food industry. More recently, biofilms are discussed as new types of living materials with tuneable mechanical properties. In particular, *Escherichia coli* produces a matrix composed of amyloid-forming curli and phosphoethanolamine-modified cellulose fibres in response to suboptimal environmental conditions. It is currently unknown how the interaction between these fibres contributes to the overall mechanical properties of the formed biofilms and if extrinsic control parameters can be utilized to manipulate these properties. Using shear rheology, we show that biofilms formed by the *E. coli* K-12 strain AR3110 stiffen by a factor of two when exposed to the trivalent metal cations Al(III) and Fe(III) while no such response is observed for the bivalent cations Zn(II) and Ca(II). Strains producing only one matrix component did not show any stiffening response to either cation or even a small softening. No stiffening response was further observed when strains producing only one type of fibre were co-cultured or simply mixed after biofilm growth. These results suggest that the *E. coli* biofilm matrix is a uniquely structured composite material when both matrix fibres are produced from the same bacterium. While the exact interaction mechanism between curli, phosphoethanolamine-modified cellulose and trivalent metal cations is currently not known, our results highlight the potential of using extrinsic parameters to understand and control the interplay between biofilm structure and mechanical properties. This will ultimately aid the development of better strategies for controlling biofilm growth.

**Table of Contents Graphic:** 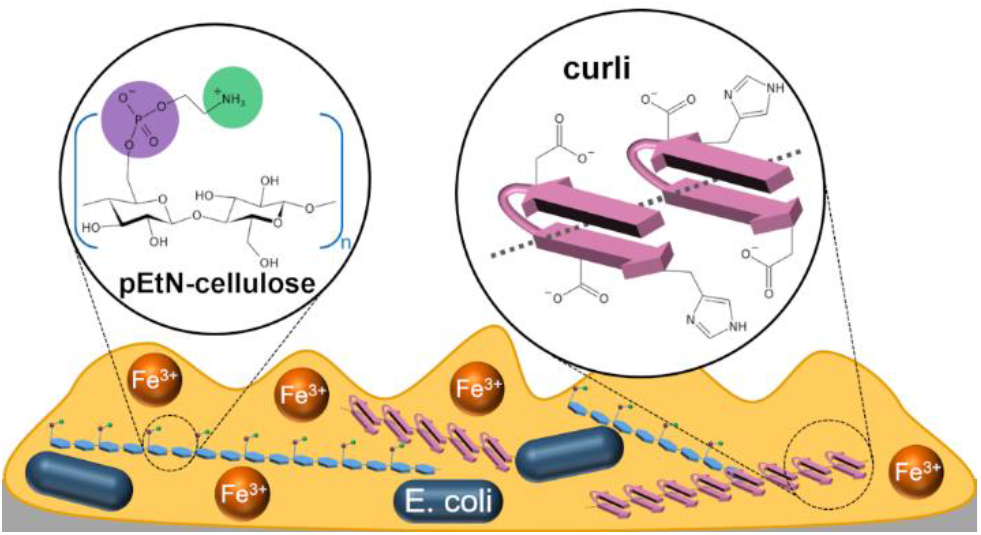

## Introduction

Biofilms are heterogeneous structures made of bacteria embedded in a self-secreted extracellular matrix. They cause complications in various fields of human life, e.g. in the medical sector,^1^ the food industry^2^ and during wastewater treatment.^3^ To date, most biofilm research has focussed on the development of preventive anti-biofilm strategies. More recently, biofilms have emerged as a potential source of sustainable materials. For example, biofilms were utilized for the formation of cement-like glue,^4^ aquaplastics^5^ or 3D-printed living materials.^6,7^ The composition of the biofilm matrix and the interaction between matrix components critically determines its mechanical properties. The matrix mainly consists of polysaccharides,^8^ proteins^9^ and nucleic acids.^10^ The type of protein and polysaccharide as well as their proportion vary remarkably, both between genera and between different species within the same genus.^11^ Protein-based amyloid fibres are particularly widespread in microbial biofilms and were, for example, observed in *Pseudomonas sp., Bacillus sp*. and *E. coli* biofilms, where they are referred to as curli fibres.^12^ Curli fibres, encoded by the *csgBA* operon, are composed of several CsgA units that polymerise onto the CsgB nucleator protein.^13^ The second main component of *E. coli* biofilms is phosphoethanolamine-modified cellulose (pEtN-cellulose).^14^ The matrix of the biofilm-forming *E. coli* K-12 strain AR3110 was estimated to contain 75 % curli and 25 % pEtN-cellulose.^15^

In addition to the matrix composition, also environmental factors influence the mechanical properties of biofilms. For instance, substrate water content, temperature, pH and nutrients may be utilized as possible control parameters for tuning *E. coli* biofilm properties.^16,17^ Another possible parameter is the addition of specific metal ions. Metal ions frequently bind to protein or carbohydrate structures in biological materials,^18^ either forming mineralized composite materials^19–22^ or sacrificial and self-healing bonds.^23,24^ Bacterial biofilms frequently occur in metallic pipes or at the surface of heavy metal containing wastewaters, suggesting a possible influence of metal ions on biofilm growth and properties. For *Enterobacter asburiae, Vitreoscilla sp*. and *Acinetobacter lwoffii*, metal ions promote biofilm formation.^25^ In the case of *Staphylococcus epidermidis, Bacillus subtilis* and *Pseudomonas aeruginosa*, biofilms stiffen in the presence of metal cations.^26^ Specifically, *B. subtilis* biofilms stiffen and erode more slowly in presence of Fe(III) and Cu(II).^27^

In the present work, we focused on *E. coli* biofilms and investigated their viscoelastic properties in the absence and presence of the bivalent metal cations Zn(II) and Ca(II) as well as the trivalent cations Al(III) and Fe(III). Performing shear rheology, we compared homogenized biofilm samples where a solution of a specific metal cation was added and samples where the same volume of water was added as a control. We further compared the *E. coli* K-12 strain AR3110, which produces biofilms with curli and pEtN-cellulose, with two closely related strains that synthesize either curli or pEtN-cellulose fibres.^14,28^ Only biofilms that contain both matrix fibres stiffen when incubated with trivalent metal ions. When strains that produce only curli or pEtN-cellulose are co-cultured or simply mixed, no cation-induced stiffening is observed, indicating that both matrix fibres need to be produced from the same bacterial cell. These results suggest the formation of a composite material during matrix production.

## Materials and Methods

### Bacterial strains

Three different *E. coli* strains were used to distinguish between the contributions of the two main matrix fibres to the mechanical biofilm properties and the dependence of these properties on the presence of metal cations. W3110 is a non-pathogenic K-12 strain^29^ that produces curli amyloid fibres and lacks the ability to synthesize cellulose. Cellulose synthesis, which is encoded in the *bcs* operon, was restored in the strain AR3110.^28^ This W3110-based strain thus produces both curli amyloid fibres and pEtN-cellulose. To obtain a strain that produces only pEtN-cellulose, curli production was inactivated in the strain AP329.^14^ To test biofilm properties when both curli and pEtN cellulose are present, but not produced by the same bacterial cell, W3110 and AP329 were combined before inoculation (co-seeded) or when harvesting the mature biofilms for the rheology experiments (mixed).

### Metal solutions

The following salts were used to probe the influence of trivalent and bivalent cations on biofilm properties: aluminium chloride hexahydrate (97%; 26726139, Molekuka GmbH), iron(III) chloride anhydrous (I/1035/50, Fisher Scientific), zinc chloride (≥98%) (29156.231, VWR International), calcium chloride dihydrate (≥99%; C3306, Sigma-Aldrich). AlCl_3_, FeCl_3_, ZnCl_2_ and CaCl_2_ were dissolved in ultrapure water to a concentration of 220 mM and the pH was measured with a pH-meter (WTW GmbH; Table 1). Using the FeCl_3_ solution as a reference, a control solution with identical pH was prepared with hydrochloric acid (1.09057, Merck KGaA). In addition to the pH, the osmolality of the metal solutions can also influence biofilm properties *via* water intake of the biofilm. The osmolalities of the different solutions were measured with an osmometer (Osmomat 3000, Gonotec GmbH). The osmolalities were determined from a calibration curve established from solutions of sodium chloride (39781.02, Serva Eletrophoresis) (Table 1, Figure S1). Similar to the pH control, a NaCl solution was prepared that matched the osmolality of the FeCl_3_ solution.

**Table 1.** Concentration, pH and osmolality of the four metal solutions FeCl_3_, AlCl_3_, ZnCl_2_, CaCl_2_ and the NaCl and HCl control solutions.

| Solution | $\text{AlCl}_3$ | $\text{FeCl}_3$ | $\text{ZnCl}_2$ | $\text{CaCl}_2$ | NaCl | HCl |
| --- | --- | --- | --- | --- | --- | --- |
| Concentration (mM) | 220 | 220 | 220 | 220 | 409 | 32 |
| pH | 2.8 | 1.5 | 5.7 | 5.2 | - | 1.5 |
| Osmolality (mOsmol/kg) | 895 | 754 | 598 | 618 | 754 | - |

### Biofilm growth

For starting the bacterial culture, LB agar plates (Luria/Miller; x969.1, Carl Roth GmbH) were prepared. A bacterial suspension, grown from glycerol stocks, was streaked onto these agar plates to obtain microcolonies after overnight culture at 37 °C. One day before starting biofilm growth, two single microcolonies were separately transferred into LB medium (5 mL; Luria/Miller; x968.2, Carl Roth GmbH) and incubated overnight at 250 rpm and 37 °C. The OD_600_ of the resulting bacteria suspensions was measured after a 10-fold dilution. The suspension where OD_600_ was closest to 0.5 was chosen for inoculating the biofilms. Biofilms were grown on salt-free LB agar plates as media with low osmolarity promote matrix production.^30^ The salt-free LB agar plates were composed of tryptone/peptone ex casein (10 g L^−1^; 8952.1, Carl Roth GmbH), yeast extract (5 g L^−1^; 2363.1, Carl Roth GmbH) and bacteriological agar agar (18 g L^−1^; 2266.3, Carl Roth GmbH). On each Petri dish (ø= 145 mm), 9 × 5 μL of suspension were inoculated to obtain an array of 9 biofilms. For the “co-seeded” biofilm samples, OD_600_ of the two suspensions was measured and the suspensions were combined such that the final density of each bacterial strain was identical. Inoculation took place immediately after a short mixing step. For the “mixed” samples, both bacterial strains were grown on the same agar surface. All biofilms were grown at 28 °C for 7 days and then stored in the fridge at 5 °C for a maximum of 48 h. Images of the biofilms were acquired with an AxioZoomV.16 stereomicroscope (Zeiss, Germany).

### Sample preparation for rheology experiments

Depending on the *E. coli* strain, two or three biofilms (^~^90 mg) were scraped from the agar surface and transferred into an empty Petri dish using cell scrapers. For the “mixed” biofilm samples, materials from both strains were combined in equal proportions. All samples were gently stirred with a pipette tip and either measured as obtained (neat) or incubated with the desired metal or control solution (diluted). For the experiments that required the incubation of the biofilm with the respective solution, the scraped biofilms were stirred with the solution in a ratio of 10:1 (w/v), yielding a final cation concentration of ^~^20 mM. After stirring, the Petri dish was sealed with Parafilm and left to incubate at room temperature for 45 min. For every dilution experiment, two samples from the same agar plate were measured. One was incubated with the solution of interest and the other sample was incubated with ultrapure water. To document sample texture, images of the different mixtures were taken with a 2 megapixel USB camera (Toolcraft Microscope Camera Digimicro 2.0 Scale, Conrad Electronic SE).

### Rheology measurements

The measurements were performed with an oscillatory shear rheometer (MCR301, Anton Paar GmbH) under stress control. The sample stage was equipped with Peltier thermoelectric cooling and the temperature was set to 21 °C for all measurements. Once the sample was transferred onto the stage, a channel around the stage was filled with water and a hood was used to maintain a high humidity environment. A parallel plate geometry (ø= 12 mm) was used and the gap was set to 250 μm.

To quantify the viscoelastic properties of the biofilm, strain amplitude sweeps were carried out to determine the linear viscoelastic range (LVE) and to extract the storage (G′_0_) and loss (G″_0_) moduli. The oscillation frequency was set to 10 rad s^−1^. The strain amplitude was increased from 0.01 % to 100 % with 7 points per decade and then decreased again. These cycles of ascending and descending strain amplitude were repeated 3x. One experiment with 3 cycles lasted approximately 45 min. The data presented in the Results section were extracted from the ascending amplitude sweep in the second cycle. The first cycle was considered as an additional homogenisation step.

To validate the chosen oscillation frequency, frequency sweeps were performed for AR3110 samples. The strain amplitude was set constant to 0.02 %. The oscillation frequency was decreased step-wise from 100 rad s^−1^ to 1 rad s^−1^ with 7 points per decade. Alternatively, frequency sweeps were also performed with a frequency increasing from 1 to 100 rad s^−1^. This ranges from one order of magnitude above and below the frequency used for the amplitude sweeps. Frequency sweeps were also performed over a wider range of frequencies, i.e. from 100 to 0.001 rad s^−1^; however, these measurements showed excessive drying of the biofilm samples at low frequencies. All frequency sweeps were carried out with neat biofilms and samples mixed with 10 % (v/w) ultrapure water, and both preceded or not by a pair of increasing and decreasing amplitude sweeps as previously described. To validate that sample drying does not affect the data acquired within the second ascending amplitude sweep, sample properties of AR3110 were recorded for a duration of at least 3 h, using a low oscillation frequency of 10 rad s^−1^ and strain amplitude of 0.02 %. This test was also preceded by a pair of amplitude sweeps (increasing and decreasing strain amplitude) as previously described.

### Data analysis

To determine biofilm properties, the G′ and G″ values were averaged over a strain range from 0.01 to 0.02 % (3 data points). These values represent the plateau moduli G′_0_ and G″_0_ of the respective biofilms (neat samples vs. samples diluted with ultrapure water). For the dilution experiments with solutions of metal cations, we primarily focussed on the relative difference between moduli. That is, the modulus of the sample diluted with the solution of interest was corrected by the modulus of a sample (from the same Petri dish) diluted with ultrapure water. This comparison to a reference sample, grown under identical conditions, was necessary to account for biofilm sample variability between Petri dishes.

For both moduli, the relative difference was calculated as follows, as exemplary shown for G′_0_:

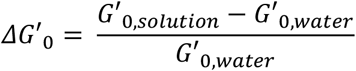

For each condition tested, the median was determined (n_pairs_ ≥ 4) as the data was not normally distributed. The data is shown in the form of boxplots. The whiskers of the boxplots represent 1.5 times the interquartile range (IQR). To assess whether the relative differences of the moduli show a significant difference from zero, i.e. the effect of the solution tested differs from that of water, a one-sample Wilcoxon signed rank test (μ= 0, a = 0.05) was performed, using the program R (R Core Team; version 4.0.5).

## Results

Biofilms that synthesize both curli and pEtN-cellulose (AR3110) showed the typical morphology with three-dimensional wrinkles (Figure 1).^14^ In contrast, the strains producing only curli (W3110) or pEtN-cellulose (AP329) showed different morphologies in agreement with the literature.^14,28^ When co-seeding W3110 and AP329, the biofilm morphology was similar to AR3110, suggesting that the structural and mechanical properties of the matrix are at least partly restored in the co-seeded biofilm.

**Figure 1.**
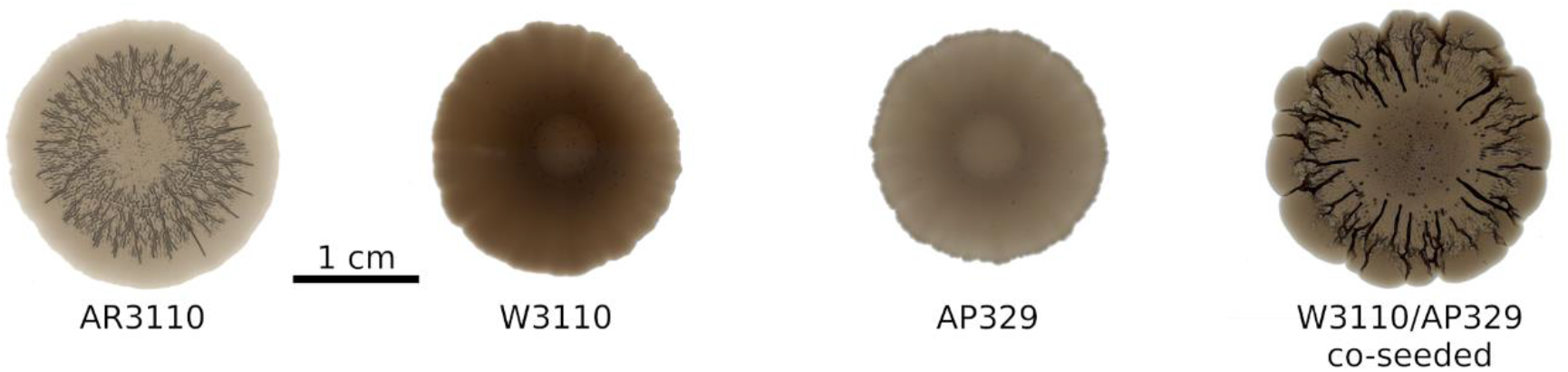
Phenotypes of the different *E. coli* strains. AR3110 produces both curli fibres and pEtN-cellulose. W3110 expresses only curli, while AP329 synthesizes only pEtN-cellulose. The sample W3110/AP329 shows the biofilm morphology obtained when W3110 and AP329 were co-seeded, i.e. when curli and pEtN-cellulose were produced by different bacteria.

For measuring the viscoelastic properties, the biofilms were harvested and mildly homogenised by stirring. It has previously been suggested that homogenized *P. aeruginosa* biofilms quickly regain their viscoelastic properties when probed with shear rheology.^31^ Here, homogenisation was necessary to mix the harvested biofilms with the metal cation solution of interest. As trivalent metal ions, Al(III) and Fe(III) were chosen for their known effects on the viscoelastic properties of *B. subtilis* and *P. aeruginosa* biofilms. Fe(III) has coordination numbers ranging from 4 to 6,^32^ Al(III) has 4 and 6, rarely 5.^33^ Zn(II) and Ca(II) were chosen as two bivalent cations with different preferred coordination numbers (Zn(II): 4-6, Ca(II): 6-8).^32,34^

### Selection of measurement conditions and data range for rheology analyses

Before probing the influence of bivalent and trivalent cations on the mechanical properties of the different biofilms, we first established the measurement conditions using neat AR3110 biofilms. As previously stated, the amplitude sweeps consisted of three cycles and the data presented were extracted from the ascending amplitude sweep in the second cycle (Figure 2A). The plateau values for both moduli were similar for the three cycles, with no systematic increase or decrease of the values between the first and subsequent cycles (Figure S2). This confirms that also homogenized *E. coli* biofilms recover their stiffness within a few minutes after yielding, similarly to what was observed for *P. aeruginosa*.^31^

**Figure 2.**
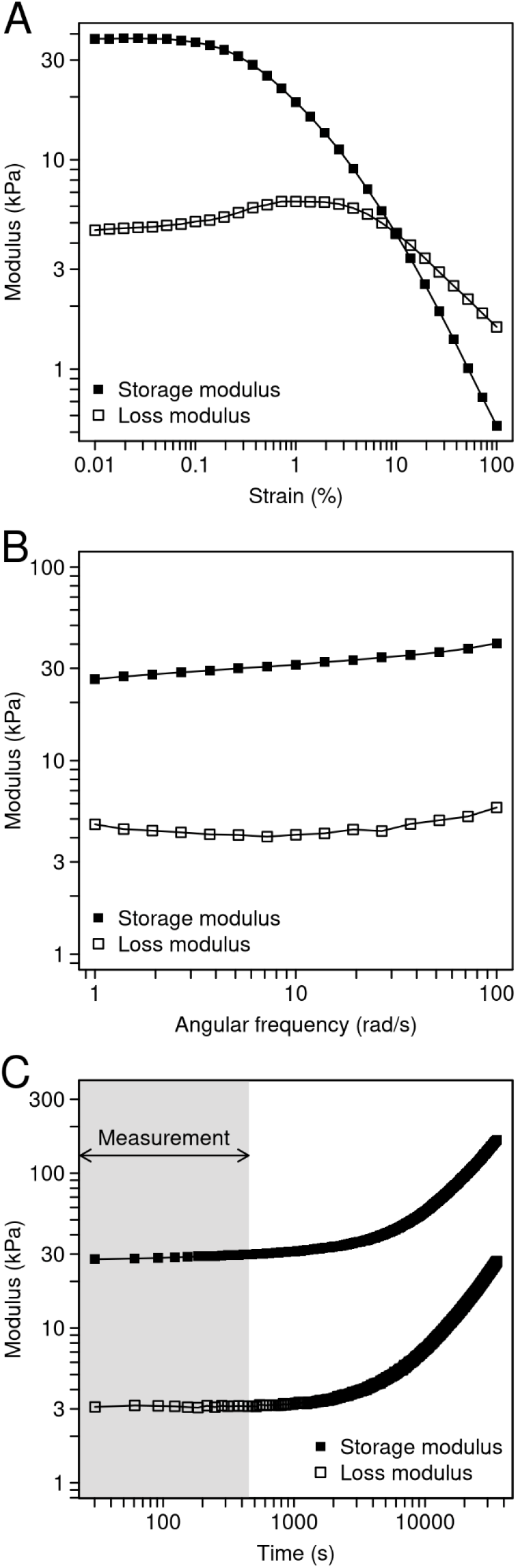
Viscoelastic properties of AR3110 biofilms, producing both matrix fibres. A) Strain amplitude sweep (ω= 10 rad s^−1^) of a biofilm where no solution was added. B) Frequency sweep (γ = 0.02 %, decreasing frequency) of a biofilm where no solution was added. C) Evolution of the storage and loss moduli, measured with constant strain amplitude (γ = 0.02 %) and frequency (ω= 10 rad s^−1^). The biofilms were measured without any solution added and the measurement was preceded by one amplitude sweep (not shown). The time interval of the analysed ascending amplitude sweep (7.5 min) is labelled in grey.

To assess the validity of the strain amplitude sweeps, frequency sweeps were performed. The storage and loss moduli showed a limited influence of the oscillation frequency over a range from 1 to 100 rad s^−1^ (Figure 2B). Similar viscoelastic properties were observed for frequency sweeps with increasing and decreasing frequency and for samples with and without the addition of 10 % (v/w) ultrapure water (Figure S3). Consequently, in the amplitude sweeps, the plateau moduli G′_0_ and G″_0_ were always obtained from the linear viscoelastic range (Figure 2A) at a frequency of 10 rad s-^1^. Frequencies below 1 rad s^−1^ were also tested but the sample showed a strong increase in the values of both moduli, supposedly due to sample drying (Figure S4).

The effect of drying was subsequently investigated in more detail. We focussed on the time window of the second ascending amplitude sweep, from which the shear moduli were derived. Although the sample appears to be continuously drying throughout the experiment, the drying effect accounts for only 10 % of the increase in G′_0_ during this period (Figure 2C). Interestingly, the values for both moduli (G′_0_ ≈ 30 kPa and G″_0_ ≈ 3 kPa) are significantly lower than those measured for the *E. coli* strain MG1655 (G′_0_ ≈100 kPa and G″_0_ ≈ 20 kPa), which produces a matrix with a different composition (curli and PGA, a linear polymer of β-1,6-N-acetyl-D-glucosamine).^35^

### Dispersion of the storage and loss moduli upon dilution

Adding metal ions in solution increases the water content of the biofilm-cation mixture. Changes in biofilm properties are thus a combined effect from the addition of water and from the respective metal ion. To disentangle these effects, we first investigated changes in biofilm viscoelasticity in response to the addition of 10 % (v/w) ultrapure water. In general, both storage and loss moduli decreased by approximately one order of magnitude. For example, for AR3110 biofilms, the storage modulus decreased from 30 to 4 kPa (Table 2) and the loss modulus was lowered from 3 to 0.4 kPa (Table 3). This indicates that the architecture of the biofilm matrix is partially destroyed when the sample is stirred after the addition of water. This observation relates to results obtained in *P. aeruginosa* biofilms where the addition of 5 % (v/w) water led to a stiffness decrease of 40 %.^31^ In most cases, the addition of water also increased the dispersion (coefficient of variation) in both moduli (Tables 2, 3, S1, S2). Considering the overall large dispersion between biofilm samples grown on different days and as a result of stirring, the following measurements to probe the effect of metal ions were performed with an internal control. Each metal containing sample was compared to a sample containing 10 % (v/w) ultrapure water that was grown in the same Petri dish (Figure 3A; Materials and Methods).

**Table 2.** Median storage moduli (G′_0_) before (-) and after (+) dilution of the biofilms with 10 % (v/w) ultrapure water (n_expriments_ ≥ 3). The G′_0_ values of all individual experiments are reported in Table S1.

| Matrix composition | Curli pEtN-cellulose |  | Curli |  | pEtN-cellulose |  | Curli pEtN-cellulose (mixed) |  | Curli pEtN-cellulose (co-seeded) |  |
| --- | --- | --- | --- | --- | --- | --- | --- | --- | --- | --- |
|  | - | + | - | + | - | + | - | + | - | + |
| Water | - | + | - | + | - | + | - | + | - | + |
| Median G'<sub>0</sub> (Pa) | 28267 | 4510 | 16533 | 1580 | 18200 | 576 | 31267 | 2673 | 51167 | 5617 |
| Median absolute deviation (MAD) (Pa) | 4567 | 863 | 6517 | 367 | 9043 | 385 | 4267 | 1250 | 3233 | 2717 |
| Coefficient of variation (MAD/median) (%) | 16 | 19 | 39 | 23 | 50 | 67 | 14 | 47 | 6 | 48 |

**Table 3.**
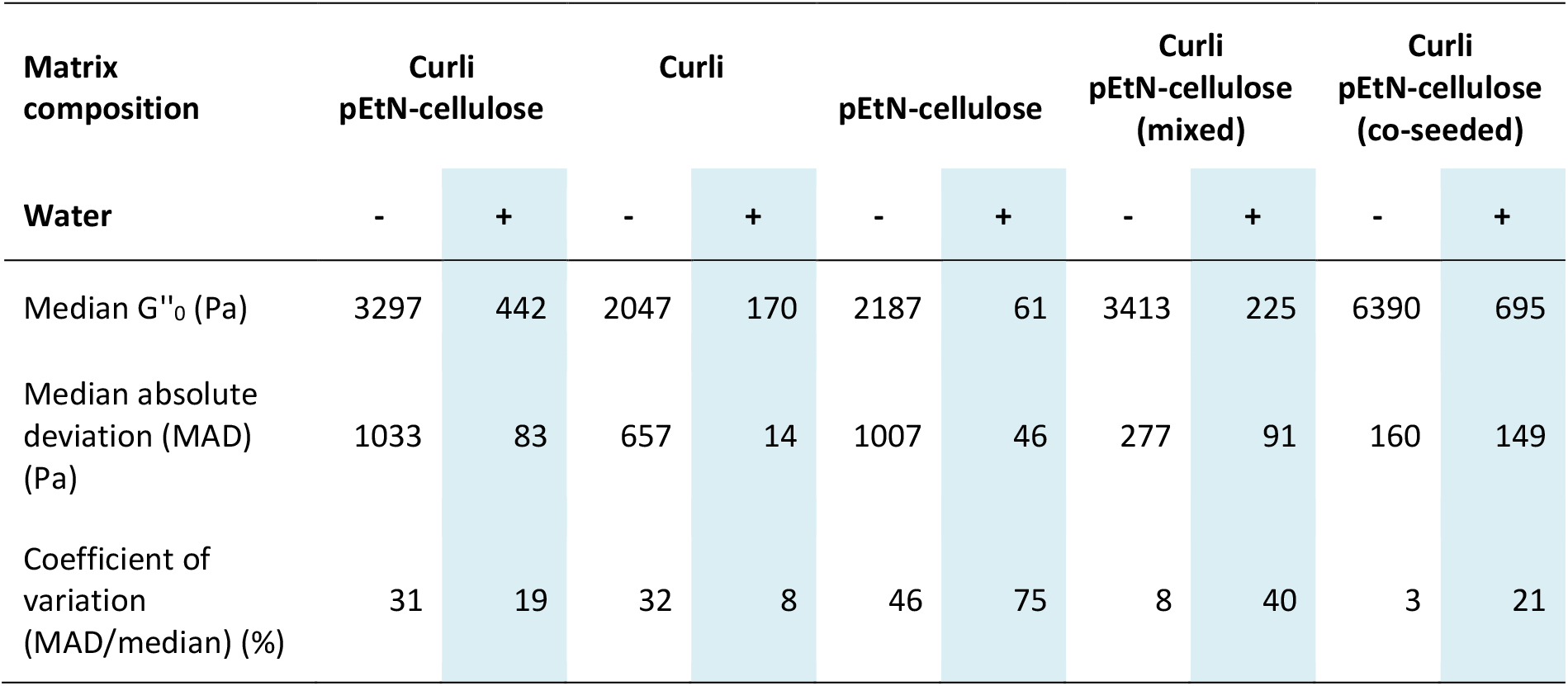
Median loss moduli (G″_0_) before (-) and after (+) dilution of the biofilms with 10 % (v/w) ultrapure water (n_expriments_ ≥ 3). The G″_0_ values of all individual experiments are reported in Table S2.

**Figure 3.**
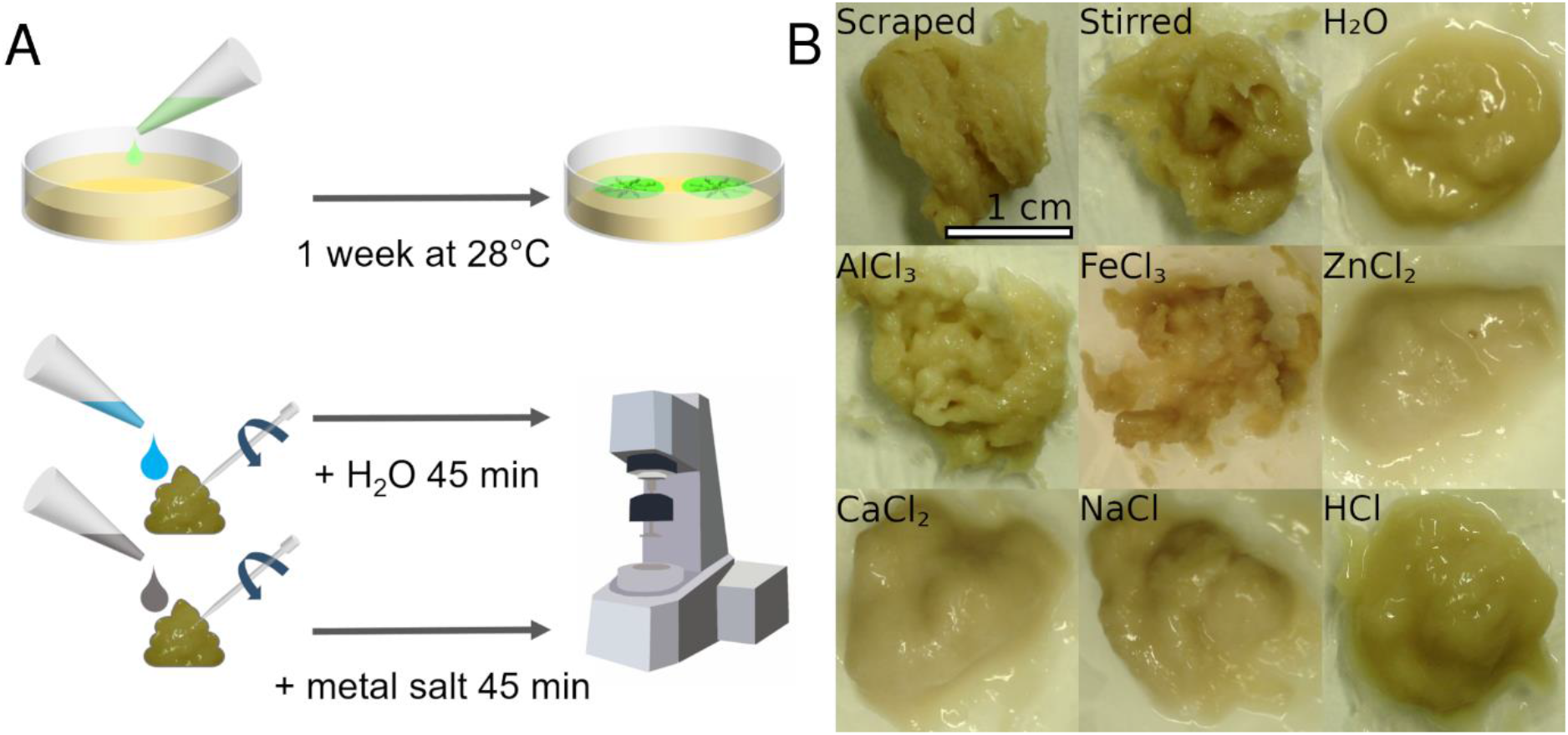
Sample preparation and texture. A) Biofilms were inoculated on salt-free LB agar and grown for one week at 28 °C. One Petri dished contained material for two rheology experiments. After harvesting two samples of the biofilm material, the metal solution of interest was added to one sample and the sample was gently stirred with a pipette tip. Ultrapure water was added to the second sample, which was then treated in the same way. Both samples were incubated for 45 min, followed by the rheology measurements. B) Texture of AR3110 biofilm material after stirring with various solutions. Biofilms stirred with ultrapure water (H_2_O), with the control solutions or solutions with bivalent metal cations appear more liquid. In contrast, the samples containing trivalent metal ions show a more solid textural appearance.

### Effect of trivalent cations on the shear modulus of AR3110 biofilms

To address the great variability between samples grown on different Petri dishes, bacteria were always seeded such that biofilm material sufficient for two samples could be obtained from the same Petri dish. After one week of growth, the biofilms were scraped from the agar. Prior to the rheology measurements, one sample was incubated with ultrapure water, while the other one was incubated with the solution of interest. This allowed a systematic comparison dish per dish between the samples incubated with a metal solution and the respective control samples incubated with water (Figure 3A).

Immediately following the addition of the metal ion solutions to AR3110 biofilms, the mixtures showed a striking difference in their visual appearance (Figure 3B). The texture of biofilms containing AlCl_3_ or FeCl_3_ was similar to a granular paste. In contrast, the biofilm mixture appeared more fluid and smooth when ZnCl_2_ or CaCl_2_ was added. No such difference between bivalent and trivalent cations was observed for any other matrix composition.

To quantify the observed texture changes, G′_0_ and G″_0_ were determined for all different biofilm samples incubated with the different metal ion solutions or the control solutions. Consistent with the changes in texture, the moduli also differed when the AR3110 biofilms were mixed with bivalent or trivalent metal cations. The addition of AlCl_3_ or FeCl_3_ increased the storage and loss moduli for AR3110 (Figure 4). Neither the bivalent metal ions caused an increase in either modulus nor did the control solutions that mimicked the pH value or osmolality of the FeCl_3_ solution (Figure 4).

**Figure 4.**
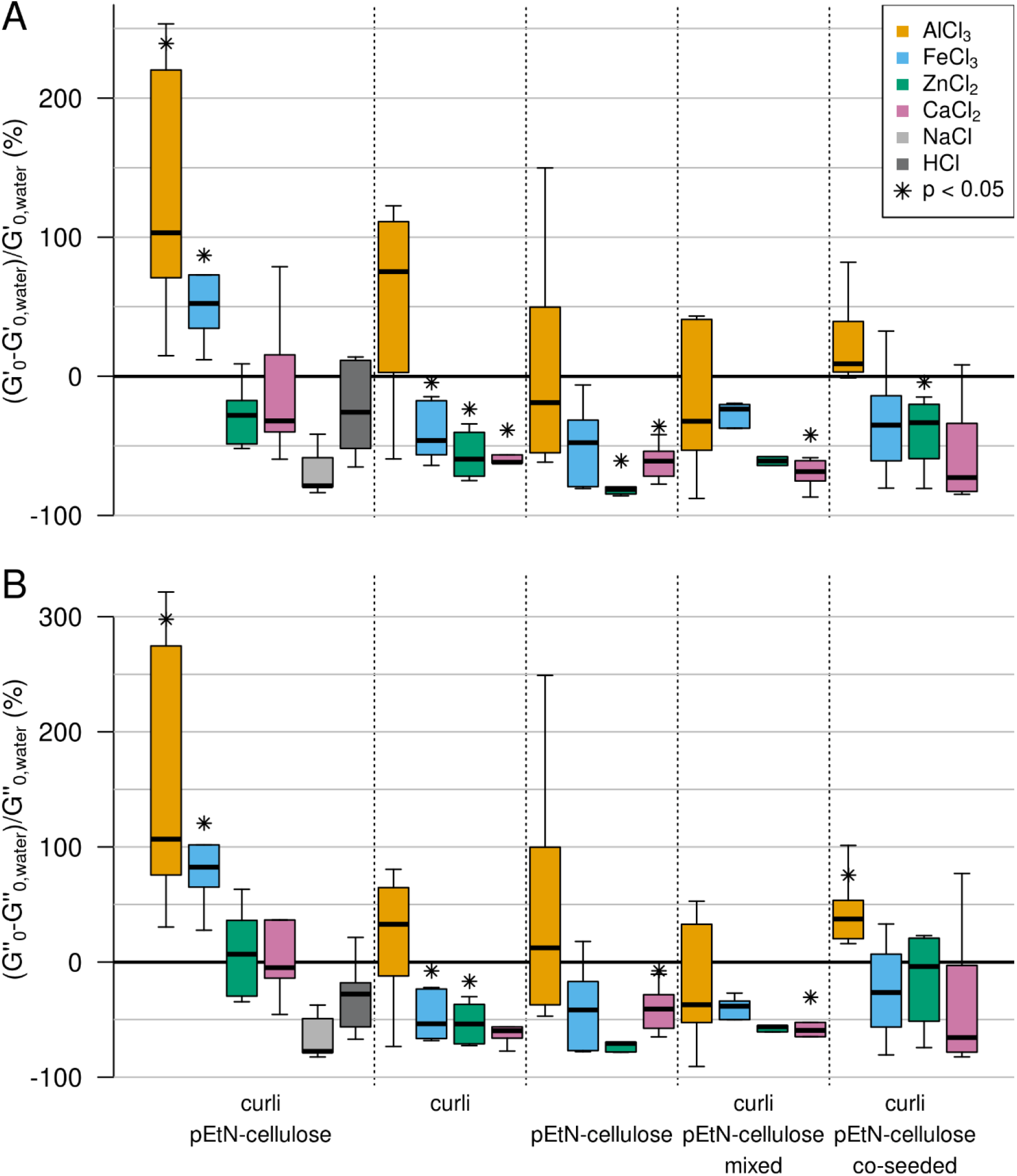
Effect of bivalent and trivalent metal ions on *E. coli* biofilms with different matrix composition. A) Storage moduli. B) Loss moduli. All samples were stirred with the respective metal cation or control solution, adding 10 % (v/w) of the respective solution. The box plots highlight the median of ≥4 independent experiments (see Tables S3-S12 for all values). The whiskers represent 1.5 times the interquartile range (IQR).

Although statistically significant (Tables 4 and 5), the increase in stiffness (G′_0_) was smaller than what was observed for other bacteria species. For example, Fe(III) and Al(III) lead to a 100-fold increase of the storage modulus of *P. aeruginosa* biofilms.^31^ Moreover, a range of bivalent and trivalent metal cations increased the storage modulus of *B. subtilis* biofilms by several orders of magnitude. Such discrepancies in the magnitude of the observed stiffening might be due to the differences in sample preparation and in matrix composition. Indeed, in the case of *P. aeruginosa*, only 5 % (v/w) solution was added,^31^ i.e. less than in our case (10 %). In *B. subtilis*, the final metal concentration in the biofilm was 0.25 M,^36^ whereas it was 0.02 M in our case. Moreover, the biofilm matrix of the *P. aeruginosa* PAO1 strain contains at least three polysaccharides (alginate, Psl, and Pel)^37^ and the *B. subtilis* B-1 strain produces mainly γ-polyglutamate, which both differ from the curli and pEtN-cellulose found in the *E. coli* biofilm matrix.

**Table 4.**
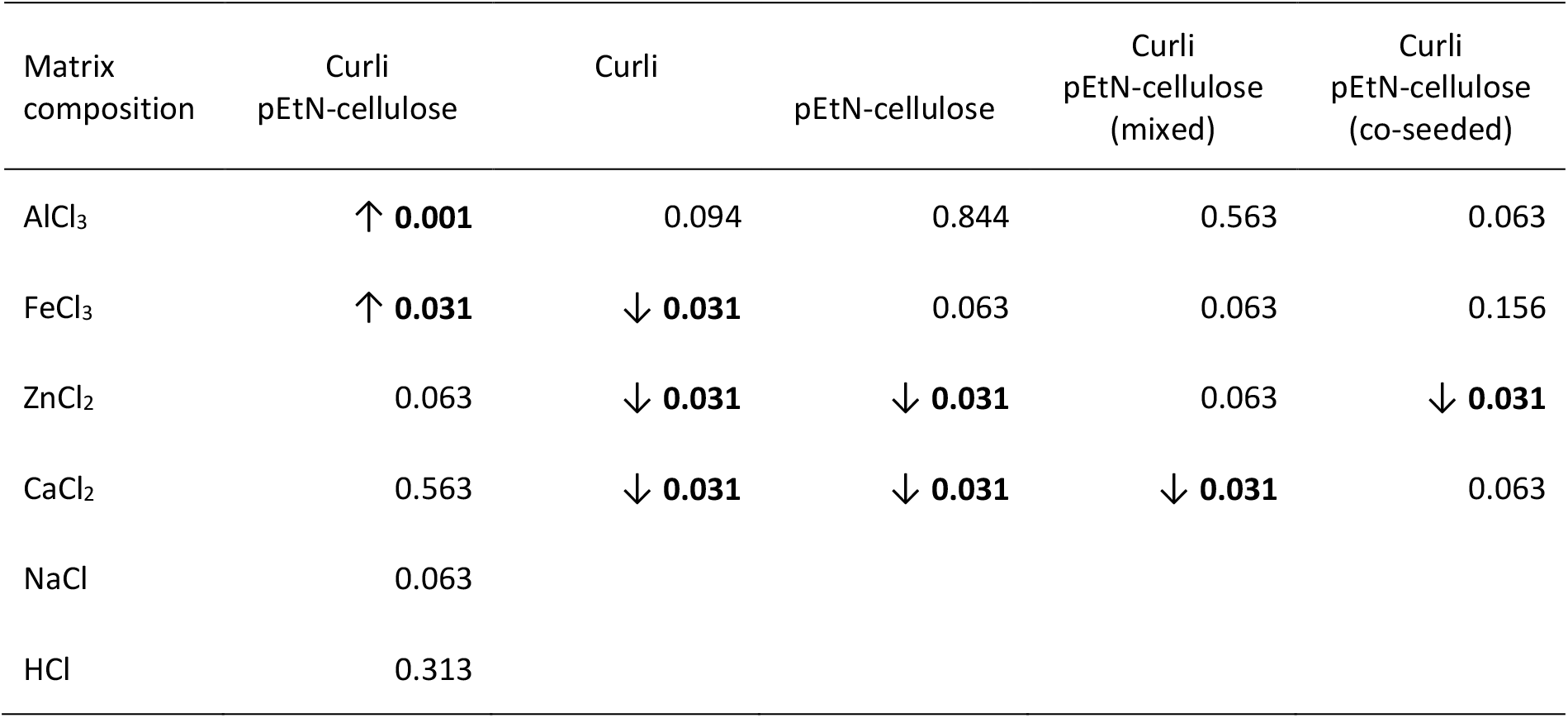
Statistical significance between the effect of a metal solution on the storage modulus (G′) and the effect of water. Shown are the p-values calculated from a One-Sample Wilcoxon Signed Rank Test (μ= 0) assessing the effect of one solution on G′ for each matrix composition. H_0_: the variation of G′ does not differ significantly from zero. p-values inferior to 0.05 in bold, in which case an arrow indicates whether the modulus increases (↑) or decreases (↓).

**Table 5.**
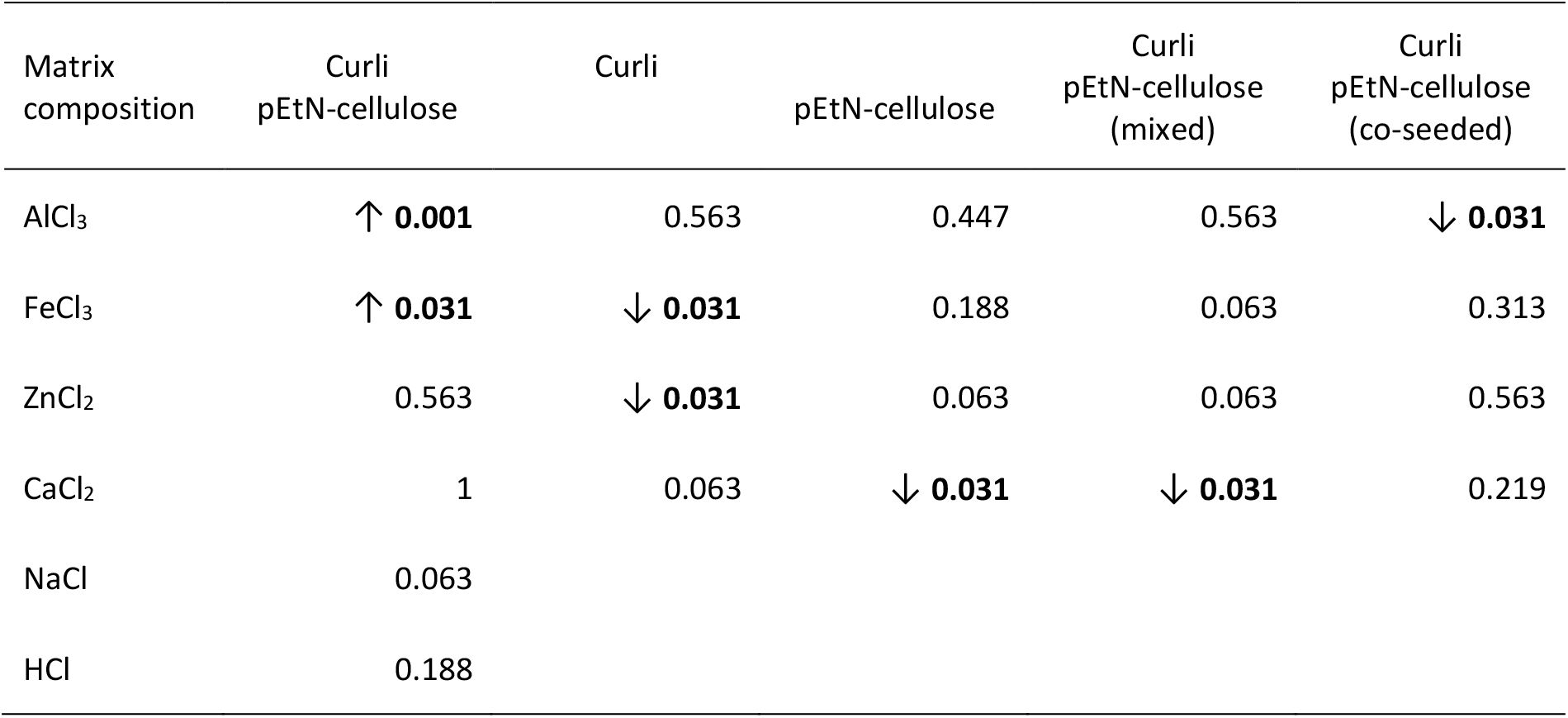
Statistical significance between the effect of a solution on the loss modulus (G″) and the effect of water. Shown are the p-values calculated from a One-Sample Wilcoxon Signed Rank Tests (μ= 0) assessing the effect of one solution on G′ for each matrix composition. H_0_: the variation of G″ does not differ significantly from zero. p-values inferior to 0.05 in bold, in which case an arrow indicates whether the modulus increases (↑) or decreases (↓).

The effect induced by Fe(III) also depends on the matrix composition (Figure 4). While the ferric salt caused a stiffening of the biofilm sample containing curli and pEtN-cellulose fibres (+50 % in G′_0_), it caused a softening (−50 % in G′_0_) for the matrix composed of curli fibres only and no statistically significant effect for the matrix composed of pEtN-cellulose. The effect remained unclear for the co-seeded and mixed biofilms. The bivalent ions caused a significant decrease (>50 %) in G′_0_ for the matrices containing only one type of fibre (Figure 4) while no such effect was observed for the AR3110 strain producing both fibres. One possible explanation for the decrease in stiffness observed for most matrix-metal combinations is a non-specific osmotic effect caused by the addition of the ionic solution. The Fe(III) - and Al(III)-induced net stiffening of the AR3110 matrix overrules this softening observed in all other samples. This suggests that the curli and pEtN cellulose fibres co-produced by AR3110 bacteria form a composite material with a built-in response to trivalent ions.

## Discussion

Using shear rheology, we examined how the viscoelastic properties of *E. coli* biofilms vary under the influence of metal cations. We probed biofilms formed by different *E. coli* strains that produce pEtN-cellulose and/or curli fibres. While the shear modulus generally decreased in the presence of metal solutions, it specifically increased when trivalent cations were added to a biofilm made from bacteria that co-produced both fibres. Metal cations trigger the formation of biofilms in *Enterobacter asburiae, Vitreoscilla sp*. and *Acinetobacter lwoffii*.^25^ Moreover, biofilms produced by *Bacillus subtilis, Pseudomonas putida* and *Shewanella oneidensis* allow for the biosorption of metal ions.^38^ In *E. coli* biofilms, the greatest biosorption performance was observed for Fe(III) when compared to Cd(II), Ni(II) or Cr(VI) but biofilm mechanical properties were not investigated.^39^ In other species, changes in mechanical biofilm properties were observed,^26^ revealing that the same ion can have opposite effects in different bacterial species. While Cu(II) reinforces *B. subtilis* B-1 biofilms, it weakens those produced by *P. aeruginosa*.^31,36^ This suggests a specific interplay between matrix composition and the type of ion. In a strain of *B. subtilis* producing a multi-component matrix, however, the effect of metal cations on the biofilm viscoelastic properties did not seem to be dictated by any specific matrix component.^27^ To interpret the present results, a molecular understanding of the possible interaction of trivalent cations with the matrix fibres is required. To our knowledge, no data is available concerning the interaction of Al(III) or Fe(III) with pEtN-cellulose. Yet, it was demonstrated that phosphorylation of cellulose nanofibers significantly enhances their adsorption capacity of Fe(III) ions.^40^ Most interestingly, phosphorylated bacterial cellulose has a much stronger affinity for Fe(III) ions than for Zn(II), in particular in acidic solutions.^41^ It was also shown that Fe(III) ions exhibit tetrahedral coordination when bound to hydroxyethyl cellulose or carboxymethyl cellulose.^42^ Tetrahedral coordination is the second most common geometry for Fe(III) after octahedral, but it is also the most common coordination geometry for Zn(II).^32^ This may suggest that the overall charge is more important than the coordination geometry.

Equally little information is available about the interaction between metal cations and amyloid curli fibres. It was demonstrated that curli fibres sequester Hg(II) ions, suggesting a possible general ability to bind metal cations.^43^ More broadly, the interaction between metal cations and other amyloid-forming structures was widely investigated. This includes amyloid beta (Aβ) peptides, which are the main components of amyloid plaques responsible for Alzheimer’s disease. While Fe(III), Al(III) and Zn(II) co-localise with Aβ in senile plaques, their influence on the *in vitro* formation of amyloid fibrils differs. Zn(II) inhibits the formation of β-sheets while both trivalent cations trigger or stabilise them.^44^ 3D-models have shown that Al(III) is almost always hexacoordinated and interacts with aspartate and glutamate residues in Aβ-complexes.^45^ Zn(II) coordinates four to six ligands in Aβ-complexes, including three histidines as well as one aspartate and/or glutamate residue.^46^ Although there is a lack of structural studies on Fe(III)-coordination to Aβ,^47^ ferric ions bind histidine more efficiently than Zn(II).^48^ A 3D-structure prediction of the major curlin subunit CsgA (AlphaFold; Figure 5) reveals close proximity of several surface-exposed histidine, glutamate and aspartate residues, suggesting that several residues are available for metal coordination.^49,50^

**Figure 5.**
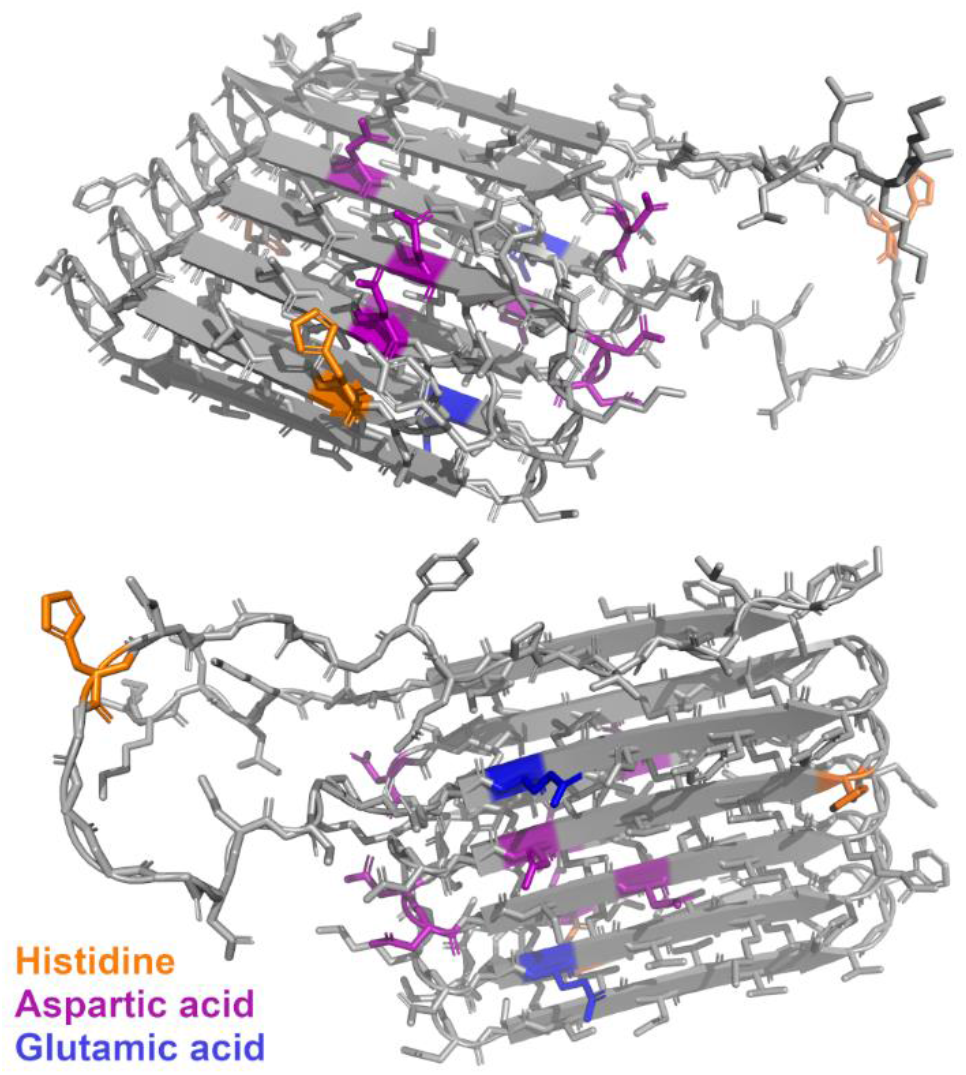
Tertiary structure of CsgA, the major curlin subunit, as predicted by AlphaFold (top and bottom view). Amino acids known to be involved in metal coordination are highlighted as follows: orange - histidine, purple - aspartic acid, blue - glutamic acid.

While our results point towards a determining role of the matrix, they do not allow us to exclude a possible effect of the metal ions on the bacteria themselves. Since metal ions trigger biofilm formation in various species,^25^ matrix production may be regulated by the presence of metal ions. Considering the timescale of our experiments, altered expression of matrix components is considered to play a minor role, however. Bacteria may further respond to reduce a possible toxic effect of heavy metal ions. For planktonic *E. coli* cells, it was shown that Fe(III) and Al(III) in a concentration of 0.01 mM reduce the number of colony forming units by 50 %.^51^ While it appears likely that biofilms provide protection against heavy metal toxicity, as demonstrated for *P. aeruginosa*,^52^ it cannot be fully neglected that these cations also have an effect on *E. coli* cells in biofilms. It is reasonable to assume, however, that the viscoelastic biofilm properties are not significantly altered by the appearance of non-viable bacteria as bacterial cells can most likely be considered as particles in a composite material. Most importantly, toxicity would affect all strains equally while we observe a clear difference between strains producing different matrix fibres.

## Conclusions

Investigating the influence of Fe(III), Al(III), Zn(II) and Ca(II) on the viscoelastic properties of *E. coli* biofilms, we observed a slight stiffening in the presence of trivalent cations. This stiffening only occurred for the strain that produced a matrix composed of both pEtN-cellulose and curli amyloid fibres. Derivatives of bacterial cellulose as well as amyloid-forming structures are known to bind metal cations; however, no molecular level information is currently available about the interaction of *E. coli* produced pEtN-cellulose and curli fibres. Considering that stiffening only occurs when both fibres are co-produced by one and the same bacterial cell, it is highly likely that the trivalent cations simultaneously interact with both components. Further research is required to unravel the molecular interactions that underlie this highly selective and specific biofilm stiffening. Towards this goal, experiments with purified and/or synthetic matrix components may provide mechanistic insights into the cation-matrix interaction. These experiments will further allow for probing the role of the phosphoethanolamine modification. Ultimately, the present work and the proposed follow-up studies will pave the way for new strategies to control biofilm viscoelastic properties without the need for genetic engineering, a topic of interest for both biofilm prevention and biofilm-based materials engineering.

## Acknowledgments

The authors thank Prof. Dr. Regine Hengge and Dr. Diego O. Serra for kindly providing the *E. coli* strains used in this work, Christine Pilz-Allen and Reinhild Dünnebacke for technical support in the laboratories, Dr. Angelo Valleriani for help with statistics, Geonho Song with rheology, Ricardo Ziege with microscopy and Wenbo Zhang with osmometry. We further thank the International Max Planck Research School (IMPRS) on Multi-Scale Biosystems for funding and support as well as Peter Fratzl and the members of the Biofilm-based Materials and the Mechano(bio)chemistry research groups (Max Planck Institute of Colloids and Interfaces) for inspiring discussions.

