## Supporting Information for "Influence of metal cations on the viscoelastic properties of *Escherichia coli* biofilms"

1. **Osmolality measurement**

NaCl solutions, spanning the concentration range from 200-550 mM, were prepared as three independent replicates for each point. The osmolalities of these standard solutions were measured, a linear regression was fitted using the least squares method with the *lm* function in R (R Core Team; version 4.0.5) and the resulting calibration curve was plotted. Subsequently, the osmolality of all used salt solutions (220 mM) was determined from three aliquots of the same solution. For the osmolality control solution used for biofilm experiments, a NaCl solution was prepared with a concentration (409 mM) that matched the osmolality of the 220 mM $FeCl_3$ solution of 754 mOsm $kg^{-1}$ (Figure S1).

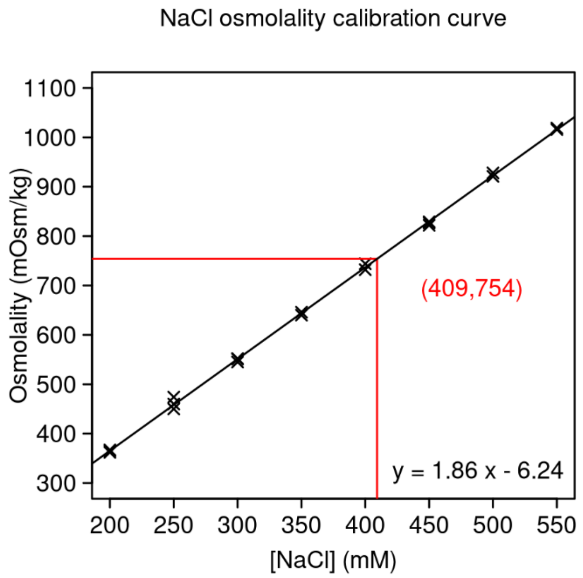

**Figure S1.** Osmolality calibration curve obtained with NaCl solutions of different concentrations. The 220 mM $FeCl_3$ solution has an osmolality of 754 mOsm $kg^{-1}$. This corresponds to a NaCl concentration of 409 mM.

2. **Amplitude sweeps for AR3110 samples with ascending and descending strain amplitude**

Amplitude sweeps were performed for AR3110 biofilms with and without the addition of 10 % (v/w) ultrapure water (Figure S2). The oscillation frequency was set to 10 rad $s^{-1}$. The strain amplitude was increased from 0.01 % to 100 % with 7 points per decade and then decreased again. These cycles of ascending and descending strain amplitude were repeated 3x. One experiment with 3 cycles lasted approximately 45 min.

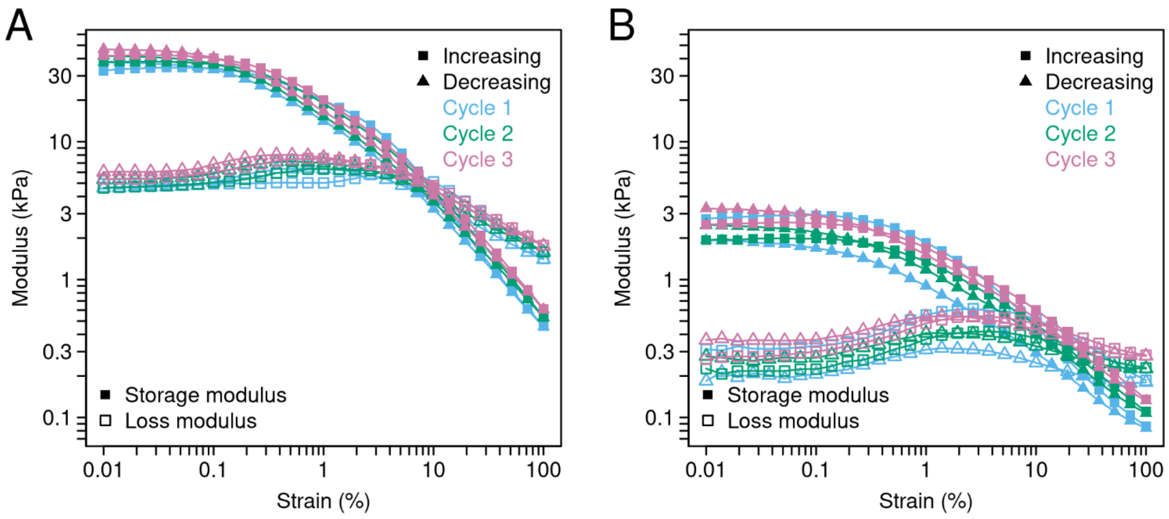

**Figure S2.** Amplitude sweeps of the strain AR3110. A) Amplitude sweep of a neat sample, consisting of three cycles of ascending and descending strain amplitudes. B) Amplitude sweep of a sample diluted with 10 % (v/w) ultrapure water, consisting of three cycles of ascending and descending strain amplitudes. The amplitude sweeps were performed at a frequency of 10 rad $s^{-1}$.

### 3. Frequency sweeps for AR3110 samples prepared under different conditions

Frequency sweeps were performed for AR3110 samples (1) with and without the addition of 10 % (v/w) ultrapure water, (2) preceded or not by an amplitude sweep or (3) starting from the highest (descending) or the lowest (ascending) frequency. The applied strain ($\gamma$ = 0.02 %) was determined from the linear viscoelastic range of the amplitude sweeps (Figure S2). The frequency range of two decades was centred on the frequency used for the amplitude sweeps ($\omega$ = 10 rad $s^{-1}$) (Figure S3).

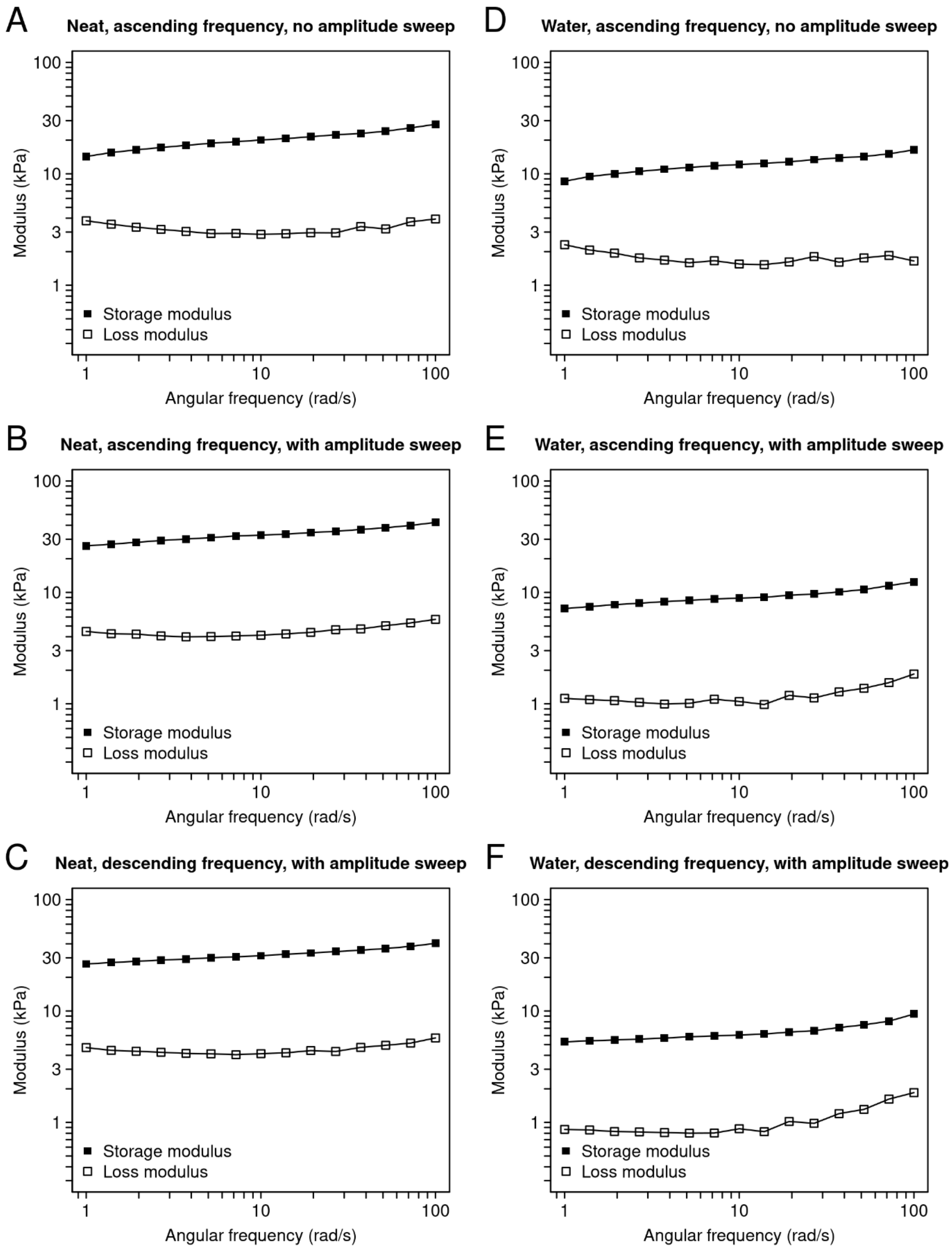

**Figure S3.** Frequency sweeps of the strain AR3110. A) Frequency sweep of a neat sample, performed with ascending frequency and γ = 0.02 %). B) Frequency sweep of a neat sample, performed with ascending frequency and γ = 0.02 %. Before the frequency sweep, the sample was subjected to one cycle of ascending and descending strain amplitudes. C) Frequency sweep of a neat sample, performed with descending frequency and γ = 0.02 %. Before the frequency sweep, the sample was subjected to one cycle of ascending and descending strain amplitudes. D) Frequency sweep of a sample diluted with 10 % (v/w) ultrapure water. The frequency sweep was performed with ascending frequency and γ = 0.02 %. E) Frequency sweep of a sample diluted with 10 % (v/w) ultrapure water. The frequency sweep was performed with ascending frequency and γ = 0.02 %. Before the frequency sweep, the sample was subjected to one cycle of ascending and descending strain amplitudes. F) Frequency sweep of a sample diluted with 10 % (v/w) ultrapure water. The frequency sweep was performed with descending frequency and γ = 0.02 %. Before the frequency sweep, the sample was subjected to one cycle of ascending and descending strain amplitudes.

In addition, a frequency sweep was performed over five decades (ω = 0.001-100 rad $s^{-1}$), using a neat or diluted sample (with preceding amplitude sweep and descending strain amplitude). The measurement was performed to determine the relaxation time of the biofilm samples; however, drying of the samples significantly affected the data in the low frequency range (Figure S4).

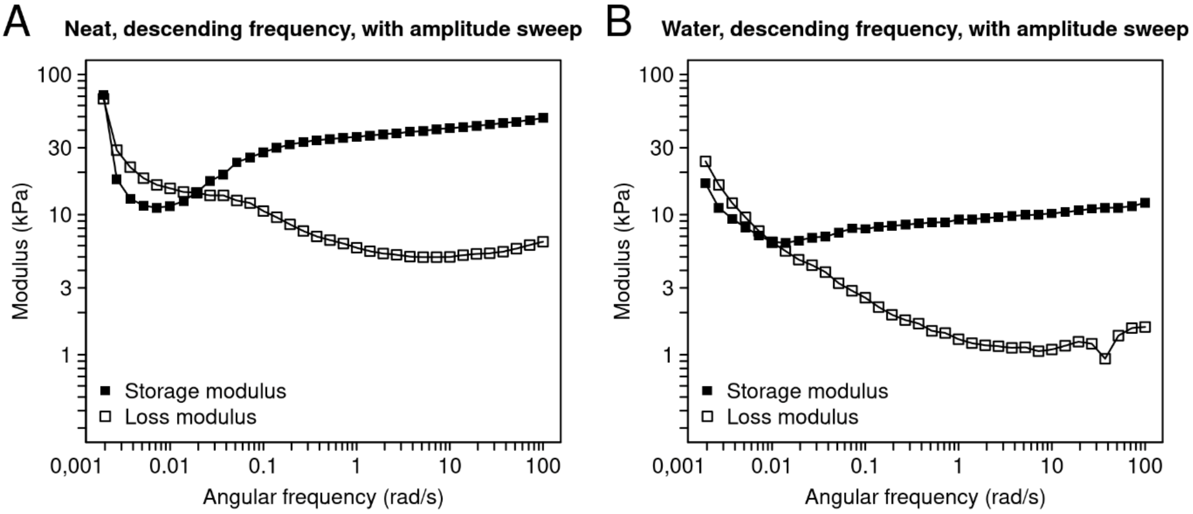

**Figure S4.** Extended frequency sweeps of the strain AR3110. A) Frequency sweep of a neat sample, performed with descending frequency and γ = 0.02 %. Before the frequency sweep, the sample was subjected to one cycle of ascending and descending strain amplitudes. B) Frequency sweep of a sample diluted with 10 % (v/w) ultrapure water. The frequency sweep was performed with descending frequency and γ = 0.02 %. Before the frequency sweep, the sample was subjected to one cycle of ascending and descending strain amplitudes.

### 4. Effect of diluting biofilm samples with ultrapure water

To determine the effect of adding 10 % (v/w) ultrapure water to the different biofilm samples, storage and loss moduli were extracted from the amplitude sweeps of ≥3 independent biofilm samples (see Methods). The extracted values, as well as their median values, are reported in Tables S1 and S2.

**Table S1.** Storage moduli ($G'_0$) of neat biofilms (-) and biofilms diluted (+) with 10 % (v/w) water.

| Matrix composition | Curli pEtN-cellulose | | Curli | | pEtN-cellulose | | Curli pEtN-cellulose (mixed) | | Curli pEtN-cellulose (co-seeded) | |
|---|---|---|---|---|---|---|---|---|---|---|
| **Water** | **-** | **+** | **-** | **+** | **-** | **+** | **-** | **+** | **-** | **+** |
| $G'_0$ (Pa) | 37933 | 1940 | 27500 | 2703 | 18200 | 576 | 27000 | 1423 | 54400 | 8333 |
| | 24400 | 2933 | 10017 | 1213 | 9157 | 190 | 31267 | 2673 | 24067 | 2857 |
| | 28267 | 5373 | 16533 | 1580 | 30567 | 1057 | 45200 | 6797 | 51167 | 5617 |
| | 32833 | 4510 | | | | | | | | |
| | 20367 | 4613 | | | | | | | | |
| **Median** | **28267** | **4510** | **16533** | **1580** | **18200** | **576** | **31267** | **2673** | **51167** | **5617** |

**Table S2.** Loss moduli ($G''_0$) of neat biofilms (-) and biofilms diluted (+) with 10 % (v/w) water.

| Matrix composition | Curli pEtN-cellulose | | Curli | | pEtN-cellulose | | Curli pEtN-cellulose (mixed) | | Curli pEtN-cellulose (co-seeded) | |
|---|---|---|---|---|---|---|---|---|---|---|
| **Water** | **-** | **+** | **-** | **+** | **-** | **+** | **-** | **+** | **-** | **+** |
| $G''_0$ (Pa) | 4663 | 217 | 4360 | 399 | 2187 | 61 | 3413 | 134 | 6390 | 843 |
| | 3197 | 359 | 1390 | 156 | 1180 | 11 | 3137 | 225 | 3430 | 367 |
| | 3297 | 539 | 2047 | 170 | 3367 | 107 | 4923 | 654 | 6550 | 695 |
| | 4330 | 479 | | | | | | | | |
| | 1950 | 442 | | | | | | | | |
| **Median** | **3297** | **442** | **2047** | **170** | **2187** | **61** | **3413** | **225** | **6390** | **695** |

### 5. Dilution experiments with metal cation solutions

To determine the effect of adding solutions of metal cations to the different biofilm samples, storage and loss moduli were extracted from the amplitude sweeps of ≥5 independent biofilm samples (see Methods). The extracted values, as well as their median values, are reported in Tables S3 to S12.

**Table S3.** Storage moduli (G'$_0$) of **AR3110** biofilms incubated with 10 % (v/w) water or a solution of interest. Pairs of values located in the same line correspond to biofilms grown in the same Petri dish. Samples incubated with NaCl and HCl were grown in the same Petri dish, which is why they share the same water control values.

| **Matrix composition** | **Curli pEtN-cellulose** | | | | | | | | | | | |
|---|---|---|---|---|---|---|---|---|---|---|---|---|
| **Solution** | **$AlCl_3$** | **Water** | **$FeCl_3$** | **Water** | **$ZnCl_2$** | **Water** | **$CaCl_2$** | **Water** | **NaCl** | **Water** | **HCl** | **Water** |
| | 7100 | 3913 | 40100 | 23200 | 7097 | 14733 | 12067 | 6750 | 3997 | 9623 | 10733 | 9623 |
| | 13333 | 4893 | 38400 | 34333 | 11367 | 13767 | 12700 | 19167 | 1803 | 11000 | 8163 | 11000 |
| | 7520 | 2270 | 33133 | 24633 | 14267 | 20500 | 8800 | 21767 | 2603 | 12633 | 4400 | 12633 |
| | 5543 | 3460 | 32700 | 21400 | 11000 | 14800 | 11367 | 18933 | 608 | 1042 | 1187 | 1042 |
| | 6797 | 4927 | 38500 | 25367 | 8553 | 16633 | 16033 | 23033 | 2023 | 9537 | 4600 | 9537 |
| G'$_0$ (Pa) | 3193 | 2780 | 171000 | 52767 | 17467 | 16033 | 23733 | 20567 | | | | |
| | 117667 | 19133 | | | | | | | | | | |
| | 14567 | 4713 | | | | | | | | | | |
| | 20600 | 5830 | | | | | | | | | | |
| | 8073 | 3973 | | | | | | | | | | |
| | 11033 | 6050 | | | | | | | | | | |
| **Q1** | **6948** | **3687** | **34450** | **23558** | **9165** | **14750** | **11542** | **18992** | **1803** | **9537** | **4400** | **9537** |
| **Q2 (median)** | **8073** | **4713** | **38450** | **25000** | **11183** | **15417** | **12383** | **19867** | **2023** | **9623** | **4600** | **9623** |
| **Q3** | **13950** | **5378** | **39700** | **32092** | **13542** | **16483** | **15200** | **21467** | **2603** | **11000** | **8163** | **11000** |

**Table S4.** Loss moduli (G"$_0$) for **AR3110** biofilm samples incubated with 10 % (v/w) water or a solution of interest. Pairs of values located in the same line correspond to biofilms grown in the same Petri dish. Samples incubated with NaCl and HCl were grown in the same Petri dish, which is why they share the same water control values.

| **Matrix composition** | **Curli pEtN-cellulose** | | | | | | | | | | | |
|---|---|---|---|---|---|---|---|---|---|---|---|---|
| **Solution** | **$AlCl_3$** | **Water** | **$FeCl_3$** | **Water** | **$ZnCl_2$** | **Water** | **$CaCl_2$** | **Water** | **NaCl** | **Water** | **HCl** | **Water** |
| | 1043 | 587 | 5690 | 2820 | 1457 | 2067 | 2520 | 968 | 447 | 878 | 1067 | 878 |
| | 2187 | 794 | 5613 | 4397 | 2253 | 1653 | 2250 | 2613 | 242 | 1373 | 1127 | 1373 |
| | 1270 | 333 | 4763 | 2883 | 2613 | 2483 | 1553 | 2847 | 380 | 1770 | 583 | 1770 |
| | 977 | 517 | 4650 | 2513 | 2017 | 1860 | 2053 | 2360 | 99 | 158 | 114 | 158 |
| | 1277 | 979 | 5383 | 2990 | 1630 | 2487 | 3093 | 2993 | 278 | 1240 | 542 | 1240 |
| G"$_0$ (Pa) | 572 | 329 | 22800 | 6117 | 2987 | 1830 | 4803 | 3517 | | | | |
| | 14633 | 2060 | | | | | | | | | | |
| | 2073 | 563 | | | | | | | | | | |
| | 2850 | 676 | | | | | | | | | | |
| | 1203 | 582 | | | | | | | | | | |
| | 1760 | 1227 | | | | | | | | | | |
| **Q1** | **1123** | **540** | **4918** | **2836** | **1727** | **1838** | **2103** | **2423** | **242** | **878** | **542** | **878** |
| **Q2 (median)** | **1277** | **587** | **5498** | **2937** | **2135** | **1963** | **2385** | **2730** | **278** | **1240** | **583** | **1240** |
| **Q3** | **2130** | **886** | **5671** | **4045** | **2523** | **2379** | **2950** | **2957** | **380** | **1373** | **1067** | **1373** |

**Table S5.** Storage moduli ($G'_0$) for **W3110** biofilms incubated with 10 % (v/w) water or a solution of interest. Pairs of values located in the same line correspond to biofilms grown in the same Petri dish.

| **Matrix composition** | **Curli** | | | | | | | |
|---|---|---|---|---|---|---|---|---|
| **Solution** | **$AlCl_3$** | **Water** | **$FeCl_3$** | **Water** | **$ZnCl_2$** | **Water** | **$CaCl_2$** | **Water** |
| $G'_0$ (Pa) | 3550 | 8737 | 3287 | 6630 | 6427 | 20967 | 3900 | 10133 |
| | 21400 | 9613 | 9393 | 11000 | 2413 | 4793 | 3737 | 8587 |
| | 11133 | 10833 | 2247 | 5153 | 2857 | 10133 | 3210 | 14667 |
| | 10500 | 4973 | 8290 | 23067 | 4717 | 7167 | 5777 | 15367 |
| | 9940 | 6103 | 3350 | 5763 | 1323 | 5283 | 4050 | 10667 |
| | 12267 | 6543 | 4947 | 6000 | 4990 | 8360 | 11333 | 11933 |
| **Q1** | **10080** | **6213** | **3302** | **5823** | **2524** | **5754** | **3777** | **10267** |
| **Q2 (median)** | **10817** | **7640** | **4148** | **6315** | **3787** | **7763** | **3975** | **11300** |
| **Q3** | **11983** | **9394** | **7454** | **9908** | **4922** | **9690** | **5345** | **13983** |

**Table S6.** Loss moduli ($G''_0$) for **W3110** biofilms incubated with 10 % (v/w) water or a solution of interest. Pairs of values located in the same line correspond to biofilms grown in the same Petri dish.

| **Matrix composition** | **Curli** | | | | | | | |
|---|---|---|---|---|---|---|---|---|
| **Solution** | **$AlCl_3$** | **Water** | **$FeCl_3$** | **Water** | **$ZnCl_2$** | **Water** | **$CaCl_2$** | **Water** |
| $G''_0$ (Pa) | 584 | 2187 | 479 | 1507 | 1353 | 3287 | 710 | 2093 |
| | 4393 | 2433 | 2030 | 2650 | 481 | 939 | 714 | 1780 |
| | 2353 | 2677 | 351 | 1042 | 516 | 1780 | 599 | 2637 |
| | 2063 | 1253 | 1630 | 3467 | 920 | 1317 | 1083 | 2477 |
| | 1833 | 1697 | 548 | 1197 | 236 | 861 | 763 | 1880 |
| | 2327 | 1477 | 888 | 1140 | 955 | 1510 | 2513 | 2097 |
| **Q1** | **1891** | **1532** | **496** | **1154** | **490** | **1034** | **711** | **1933** |
| **Q2 (median)** | **2195** | **1942** | **718** | **1352** | **718** | **1413** | **739** | **2095** |
| **Q3** | **2347** | **2372** | **1445** | **2364** | **946** | **1713** | **1003** | **2382** |

**Table S7.** Storage moduli (G'$_0$) for **AP329** biofilms incubated with 10 % (v/w) water or a solution of interest. Pairs of values located in the same line correspond to biofilms grown in the same Petri dish.

| **Matrix composition** | | | | **pEtN-cellulose** | | | | |
|---|---|---|---|---|---|---|---|---|
| **Solution** | **$AlCl_3$** | **Water** | **$FeCl_3$** | **Water** | **$ZnCl_2$** | **Water** | **$CaCl_2$** | **Water** |
| G'$_0$ (Pa) | 5337 | 3470 | 1353 | 1443 | 1977 | 3250 | 5137 | 11133 |
| | 2107 | 4933 | 1417 | 6847 | 1600 | 10367 | 4773 | 21267 |
| | 3530 | 9213 | 1623 | 8413 | 3153 | 16500 | 6960 | 18933 |
| | 9580 | 10800 | 6233 | 9103 | 2173 | 10600 | 4857 | 17233 |
| | 12533 | 17067 | 8857 | 16933 | 1203 | 8540 | 8577 | 14767 |
| | 7843 | 16567 | | | 1870 | 10233 | 8157 | 19800 |
| | 10100 | 4043 | | | | | | |
| | 10767 | 7387 | | | | | | |
| **Q1** | **4885** | **4711** | **1417** | **6847** | **1668** | **8963** | **4927** | **15383** |
| **Q2 (median)** | **8712** | **8300** | **1623** | **8413** | **1923** | **10300** | **6048** | **18083** |
| **Q3** | **10267** | **12242** | **6233** | **9103** | **2124** | **10542** | **7857** | **19583** |

**Table S8.** Loss moduli (G"$_0$) for **AP329** biofilms incubated with 10 % (v/w) water or a solution of interest. Pairs of values located in the same line correspond to biofilms grown in the same Petri dish.

| **Matrix composition** | | | | **pEtN-cellulose** | | | | |
|---|---|---|---|---|---|---|---|---|
| **Solution** | **$AlCl_3$** | **Water** | **$FeCl_3$** | **Water** | **$ZnCl_2$** | **Water** | **$CaCl_2$** | **Water** |
| G"$_0$ (Pa) | 707 | 345 | 179 | 152 | 375 | 370 | 1002 | 1397 |
| | 363 | 536 | 185 | 800 | 265 | 1220 | 933 | 2660 |
| | 571 | 1077 | 216 | 975 | 558 | 1970 | 1377 | 2450 |
| | 1913 | 1533 | 988 | 1190 | 380 | 1250 | 976 | 2297 |
| | 2450 | 2450 | 1377 | 2353 | 213 | 969 | 1800 | 2013 |
| | 1150 | 1980 | | | 334 | 1100 | 1680 | 2703 |
| | 1563 | 448 | | | | | | |
| | 1680 | 862 | | | | | | |
| **Q1** | **673** | **514** | **185** | **800** | **282** | **1002** | **983** | **2084** |
| **Q2 (median)** | **1357** | **969** | **216** | **975** | **355** | **1160** | **1189** | **2373** |
| **Q3** | **1738** | **1645** | **988** | **1190** | **379** | **1243** | **1604** | **2608** |

**Table S9.** Storage moduli ($G'_0$) for samples consisting of **mixed W3110 and AP329** biofilms incubated with 10 % (v/w) water or a solution of interest. Pairs of values located in the same line correspond to biofilms grown in the same Petri dish.

| **Matrix composition** | **Curli pEtN-cellulose (mixed)** | | | | | | | |
|---|---|---|---|---|---|---|---|---|
| **Solution** | **$AlCl_3$** | **Water** | **$FeCl_3$** | **Water** | **$ZnCl_2$** | **Water** | **$CaCl_2$** | **Water** |
| $G'_0$ (Pa) | 1950 | 15900 | 4107 | 20767 | 4153 | 9840 | 8053 | 19400 |
| | 20900 | 14833 | 17867 | 22167 | 4663 | 13000 | 7267 | 19733 |
| | 8627 | 6020 | 6640 | 10600 | 11467 | 29333 | 11800 | 47767 |
| | 6847 | 11400 | 15167 | 19833 | 8180 | 14933 | 6157 | 46500 |
| | 13167 | 28100 | 16933 | 21233 | 5023 | 36500 | 15033 | 57600 |
| | 11833 | 15700 | | | | | 11967 | 30400 |
| **Q1** | **7292** | **12258** | **6640** | **19833** | **4663** | **13000** | **7463** | **22400** |
| **Q2 (median)** | **10230** | **15267** | **15167** | **20767** | **5023** | **14933** | **9927** | **38450** |
| **Q3** | **12833** | **15850** | **16933** | **21233** | **8180** | **29333** | **11925** | **47450** |

**Table S10.** Loss moduli ($G''_0$) for samples consisting of **mixed W3110 and AP329** biofilms, incubated with 10 % (v/w) water or a solution of interest. Pairs of values located in the same line correspond to biofilms grown in the same Petri dish.

| **Matrix composition** | **Curli pEtN-cellulose (mixed)** | | | | | | | |
|---|---|---|---|---|---|---|---|---|
| **Solution** | **$AlCl_3$** | **Water** | **$FeCl_3$** | **Water** | **$ZnCl_2$** | **Water** | **$CaCl_2$** | **Water** |
| $G''_0$ (Pa) | 266 | 2863 | 582 | 3570 | 770 | 1757 | 1420 | 2983 |
| | 3170 | 2387 | 3110 | 4253 | 830 | 1850 | 1280 | 2690 |
| | 1250 | 818 | 912 | 1827 | 2130 | 5413 | 2493 | 7087 |
| | 969 | 1653 | 2360 | 3567 | 1603 | 2640 | 1060 | 6787 |
| | 2020 | 4257 | 2367 | 3840 | 870 | 6443 | 3230 | 8387 |
| | 1727 | 2567 | | | | | 2443 | 5707 |
| **Q1** | **1039** | **1837** | **912** | **3567** | **830** | **1850** | **1315** | **3664** |
| **Q2 (median)** | **1488** | **2477** | **2360** | **3570** | **870** | **2640** | **1932** | **6247** |
| **Q3** | **1947** | **2789** | **2367** | **3840** | **1603** | **5413** | **2481** | **7012** |

**Table S11.** Storage moduli (G'$_0$) for **co-seeded W3110 and AP329** biofilms incubated with 10 % (v/w) water or a solution of interest. Pairs of values located in the same line correspond to biofilms grown in the same Petri dish.

| **Matrix composition** | **Curli pEtN-cellulose (co-seeded)** | | | | | | | |
|---|---|---|---|---|---|---|---|---|
| **Solution** | **$AlCl_3$** | **Water** | **$FeCl_3$** | **Water** | **$ZnCl_2$** | **Water** | **$CaCl_2$** | **Water** |
| G'$_0$ (Pa) | 8263 | 7487 | 4540 | 9743 | 6423 | 15733 | 2890 | 19033 |
| | 16867 | 12100 | 2717 | 13833 | 8297 | 12500 | 8750 | 13233 |
| | 22300 | 20733 | 8023 | 20433 | 14500 | 17033 | 4700 | 16067 |
| | 24867 | 13667 | 25700 | 19400 | 7617 | 11367 | 5157 | 20500 |
| | 28167 | 27333 | 16200 | 18833 | 16933 | 21200 | 14867 | 13733 |
| | 19267 | 19500 | 21767 | 26167 | 5650 | 29200 | 3073 | 17900 |
| **Q1** | **17467** | **12492** | **5411** | **15083** | **6722** | **13308** | **3480** | **14317** |
| **Q2 (median)** | **20783** | **16583** | **12112** | **19117** | **7957** | **16383** | **4928** | **16983** |
| **Q3** | **24225** | **20425** | **20375** | **20175** | **12949** | **20158** | **7852** | **18750** |

**Table S12.** Loss moduli (G"$_0$) for **co-seeded W3110 and AP329** biofilms incubated with 10 % (v/w) water or a solution of interest. Pairs of values located in the same line correspond to biofilms grown in the same Petri dish.

| **Matrix composition** | **Curli pEtN-cellulose (co-seeded)** | | | | | | | |
|---|---|---|---|---|---|---|---|---|
| **Solution** | **$AlCl_3$** | **Water** | **$FeCl_3$** | **Water** | **$ZnCl_2$** | **Water** | **$CaCl_2$** | **Water** |
| G"$_0$ (Pa) | 1133 | 900 | 679 | 1277 | 1060 | 2183 | 456 | 2570 |
| | 2433 | 1583 | 327 | 1687 | 1597 | 1673 | 1697 | 1747 |
| | 3807 | 2557 | 1227 | 2817 | 2847 | 2357 | 789 | 1973 |
| | 3557 | 1767 | 3770 | 2833 | 1410 | 1457 | 826 | 2860 |
| | 4257 | 3537 | 2693 | 2520 | 3277 | 2663 | 2950 | 1667 |
| | 2877 | 2480 | 3163 | 3370 | 927 | 3600 | 472 | 2167 |
| **Q1** | **2544** | **1629** | **816** | **1895** | **1148** | **1801** | **551** | **1803** |
| **Q2 (median)** | **3217** | **2123** | **1960** | **2668** | **1503** | **2270** | **807** | **2070** |
| **Q3** | **3744** | **2537** | **3046** | **2829** | **2534** | **2587** | **1479** | **2469** |
